# Mutations associated with streptomycin resistance predicted to be highly prevalent and evolvable across bacteria

**DOI:** 10.64898/2026.06.09.731231

**Authors:** Lê Na Ngô, Andrew D. Letten, Jan Engelstädter

**Affiliations:** School of the Environment, The University of Queensland, Brisbane, Queensland 4072, Australia

## Abstract

Antimicrobial resistance is an evolutionary response to antimicrobial exposure that has been extensively studied across some bacteria, including pathogens and model organisms. Yet, for most species the capacity to develop resistance remains unresolved. Here, we used computational methods to assess patterns of streptomycin resistance evolution across the bacterial tree of life. We curated a panel of high-confidence streptomycin resistance mutations, including eight mutations in the *rpsL* gene and four mutations in the *rrs* gene. We then used this panel to screen over 20 000 bacterial genomes from diverse clades. We assessed both evolvability, defined by codon-level accessibility to resistance-conferring mutations via single-nucleotide substitutions, and intrinsic resistance, where resistance-associated variants are already present. Our results suggest that most bacterial species can readily acquire *rpsL*-resistant mutations. Furthermore, we find that approximately 7% of bacterial species intrinsically carry *rpsL* resistance variants, with a wide taxonomic distribution but notable enrichment within Alphaproteobacteria. Our study provides a global view of the streptomycin resistance mutational landscape and generates testable predictions for future research.

## Introduction

Biological populations are characterised by significant variation in their capacity to respond to natural selection. This adaptive potential, or evolvability, of a population is often quantified on the basis of the amount of adaptive genetic variation pre-existing in the population (Hansen and Pélabon 2021). However, it also includes the population’s capacity to generate such variation (Payne and Wagner 2019; Riederer et al. 2022), which is especially apparent in microbial organisms with their large population sizes and short generation times. One domain where this aspect of evolvability is particularly relevant is in understanding and tackling the evolution of antibiotic resistance.

The evolvability of antibiotic resistance is shaped by the diversity of possible resistance mutations available to a bacterial strain. For most bacterial taxa, this mutational landscape remains unresolved and/or logistically challenging to obtain (e.g., inability to culture *in vitro*). However, for some well studied taxa and antibiotics, genomic screening has provided an extensive library of known resistance mutations at the nucleotide level. This information can be leveraged to infer the accessibility of resistance mutations in unscreened taxa, allowing prediction of their resistance potential (Bonny et al. 1991; Springer et al. 2001; Sander et al. 2002). Indeed, Bolourchi et al. (2025) recently used this approach to generate predictions on the evolvability to rifampicin across the bacterial tree of life (*>*18 000 species) based on a panel of resistance mutations reported in 75 species.

This approach can also provide complementary inference on intrinsic resistance in unscreened taxa. Most antibiotics prescribed today are derived from natural microbial sources (Durand et al. 2019; Hegemann et al. 2023). Approximately two-thirds of all naturally derived antibiotics are produced by Actinomycetes, with the majority originating from the genus *Streptomyces*. These actinomycete-derived antibiotics span all important classes of drugs currently used in clinics, such as aminoglycosides (kanamycin, streptomycin), *β*-lactams (cephamycins), macrolides (erythromycin) and tetracyclines (Waksman et al. 2010; Barka et al. 2016; Hutchings et al. 2019). In addition to Actinomycetes, several other bacterial taxa, including *Bacillus* and *Pseudomonas* spp., also contribute to the natural antibiotic repertoire, producing compounds such as glycopeptides, lipopeptides, and mupirocin (Pidot et al. 2014; Hutchings et al. 2019). In microbial communities, the presence of these antibiotic producers can drive the ‘natural’ evolution of resistance mechanisms in other bacteria. Consequently, it is reasonable to expect widespread intrinsic resistance or the capacity to readily evolve resistance across diverse bacterial lineages. Systematically mapping known resistance-associated mutations across bacterial genomes provides a means to identify putatively intrinsic species, as well as taxa, with high potential to develop resistance.

In this study, we adopt the approach of Bolourchi et al. (2025) to investigate evolvability and intrinsic resistance to streptomycin (STR) across diverse bacteria. Discovered in 1944, STR is an aminoglycoside antibiotic derived from *Streptomyces griseus* (Schatz et al. 1944). It was the first antibiotic found to be effective against *Mycobacterium tuberculosis*, the causative agent of tuberculosis, and was used in control programs for many years (Bloom and Murray 1992). STR prevents protein synthesis by binding to the 16S rRNA phosphate backbone and contacting K45 in the S12 ribosomal protein. These interactions promote codon misreading, which disrupts protein synthesis and leads to bacterial cell death (Carter et al. 2000). Resistance to STR arises predominantly through mutations in the *rpsL* and *rrs* genes, encoding ribosome protein S12 and the 16S rRNA, respectively (Cuevas-Córdoba et al. 2013; Thida Oo et al. 2018; Wang et al. 2019). Together, these mutations account for approximately 70% of *M. tuberculosis* STR-resistant isolates (Sreevatsan et al. 1996; Cuevas-Córdoba et al. 2013; Jagielski et al. 2014; Smittipat et al. 2016). STR resistance due to *rpsL* resistance has also been widely reported in many other bacterial species (Timms et al. 1992; Gregory et al. 2001; Pelchovich et al. 2013; Lee et al. 2023; Schmidt et al. 2025).

To generate predictions on evolvability and intrinsic resistance to STR, we first compiled a comprehensive database of previously reported STR resistance mutations in *rpsL* and *rrs*, drawn from clinical and experimental data and diverse bacterial taxa. We then screened the published genomes of more than 20 000 bacterial species, encompassing all major phyla, to identify taxa carrying high-confidence STR resistance mutations from our panel and determine the evolvability towards STR resistance in species without those mutations. By integrating observed resistance variants with codon-level evolutionary accessibility, this work provides a phylogenetically broad view of the evolution of STR resistance in bacteria.

## Materials and methods

This work built upon a previously developed bioinformatics pipeline designed to study rifampicin resistance mutations in *rpoB* (Bolourchi et al. 2025). The majority of the codebase from that pipeline was adapted and extended here to examine streptomycin (STR) resistance, an aminoglycoside antibiotic for which resistance mutations occur in the *rpsL* and *rrs* genes.

### Selection of mutation panel

First, we performed a comprehensive review of studies reporting mutations associated with phenotypic streptomycin resistance. Data were gathered from natural or clinical isolates, and from mutants derived experimentally. We conducted a comprehensive literature search on the Web of Science using the following key terms: (streptomycin AND (resistance OR resistant) AND (*rpsL* OR “ribosomal protein” OR “S12” OR *rrs* OR “16S ribosomal RNA” OR “16S rRNA”)). The titles and abstracts were screened to identify relevant articles. We then manually examined the full text of selected publications to extract species names, origins (experimental studies vs. isolates), specific mutations, and their corresponding positions. To standardize amino acid and nucleotide positions, all sequences were aligned with the *Escherichia coli* MG1655 reference. Specifically, *rpsL* sequences were aligned with the *E. coli* MG1655 *rpsL* amino acid sequence, while *rrs* sequences were aligned with the *E. coli* MG1655 *rrnB* 16S rRNA sequence. To minimize inadvertent inclusion of spurious resistance mutations, only *rpsL* substitutions reported in at least three independent studies and in at least three species were retained. Since only a few studies have reported STR resistance mutations associated in *rrs*, we applied a less stringent filter for *rrs*, retaining only mutations reported in at least two studies and in at least two species

### Predicting the presence and evolvability of STR resistance-conferring mutations

We next retrieved all available bacterial reference genomes from the NCBI RefSeq Genome database (O’Leary et al. 2024). This encompassed genomes across all assembly levels, including contigs, scaffolds, and complete assemblies, to ensure a broad representation of genomic diversity. Subsequently, *rpsL* and *rrs* sequences were extracted from the annotated genomes. Specifically, *rpsL* sequences were identified using the keywords ‘*rpsL*’, ‘30S ribosomal subunit protein S12’ and ‘30S ribosomal protein S12’, with only sequences *≥* 300 bp retained. The *rrs* sequences were retrieved using the keywords ‘*rrs*’, ‘16S ribosomal RNA’, and ‘16S rRNA’, with sequences shorter than 1200 bp excluded.

All sequences were then aligned with the *E. coli* MG1655 reference sequences (*rpsL* and *rrnB* 16S rRNA, respectively). To enhance the robustness and reliability of the analysis, the Levenshtein distance between each sequence and its reference was computed to identify and exclude highly divergent or potentially low-quality sequences. Sequences with excessive divergence were excluded, using a threshold of *≥* 40 for *rpsL* and a more permissive cut-off of *≥* 150 for *rrs*, reflecting the greater length and variability of 16S rRNA. In addition, a minimum alignment score of *−*1000 was applied for *rrs*.

The filtered *rpsL* sequences were utilized to assess intrinsic resistance and evolvability. For each sequence–mutation combination, intrinsic resistance was determined as the presence of the reported mutation, while evolvability was quantified as the number of possible single nucleotide substitutions (including zero) that could lead to that mutation at the amino acid level. Subsequently, these results were used to calculate overall predicted resistance, defined as the presence of at least one resistance mutation, and overall evolvability, measured as the total number of resistance mutations that can evolve. As *rrs* is not codon-based and all resistance mutations are accessible via single-step changes, it was evaluated only for intrinsic resistance.

### Phylogenetic analyses

To determine the phylogenetic distribution of STR-resistant species and their evolvability, we utilized a comprehensive bacterial phylogenetic tree from the Genome Taxonomy Database (GTDB; https://gtdb.ecogenomic.org) (Parks et al. 2022). The GTDB taxonomy is based on genomic trees generated from an aligned concatenated set of 120 single-copy markers for bacteria. We extracted a subtree that contained only the species identified in our final screening. We incorporated the GTDB taxonomy for consistent labeling throughout the bacterial tree. We assessed phylogenetic clustering of resistance and evolvability by computing Blomberg’s K and Pagel’s *λ* statistics (Pagel 1999; Blomberg et al. 2003).

### Implementation

All analyses in this study were performed using R version 4.5.1 (R Core Team 2025), with all code available at https://github.com/lna0104/Streptomycin. Bioinformatics analyses were performed using packages rentrez v1.2.4 (Winter 2017), ALJEbinf v0.1.0 (Engelstädter 2025), Biostrings v2.78.0 (Pagès et al. 2024), pwalign v1.6.0 (Aboyoun and Gentleman 2024), stringr v1.6.0 (Wickham 2023), future.apply v1.20.1 (Bengtsson 2021), MSA2dist v1.14.0 (Ullrich 2024). Phylogenetic analysis and tree visualization were conducted using ape v5.8.1 (Paradis and Schliep 2019), castor v1.8.4 (Louca and Doebeli 2018), ggtree v4.0.4 (Xu et al. 2022), tidytree v0.4.7 (Yu 2017), and treeio v1.34.0 (Wang et al. 2020). Data wrangling and visualisation were performed using packages tidyverse v2.0.0 (Wickham et al. 2019), ggnewscale v0.5.2 (Campitelli 2025), GGally v2.4.0 (Schloerke et al. 2024), ggpubr v0.6.2 (Kassambara 2023), ggh4x v0.3.1 (Brand 2024), tidygraph v1.3.1 (Pedersen 2024c), ggraph v2.2.2 (Pedersen 2024a), igraph v2.2.1 (Csárdi et al. 2025), RColorBrewer v1.1.3 (Neuwirth 2022), wesanderson v0.3.7 (Ram and Wickham 2023), patchwork v1.3.2 (Pedersen 2024b), and cowplot v1.2.0 (Wilke 2024). For interactive protein visualisation, the NGLVieweR v1.4.0 package (Velden 2024) was used in combination with htmlwidgets v1.6.4 (Vaidyanathan et al. 2023).

## Results

### Mutations conferring STR resistance

We compiled data from 98 studies across 40 species that reported a total of 311 mutations in the *rpsL* gene associated with STR resistance. After merging homologous residues reported across different studies and species, we identified 66 unique single amino acid substitutions at 31 positions. These mutations are distributed across both laboratory-generated variants and those isolated from clinical or environmental samples. To identify high-confidence mutations, we implemented stringent selection criteria, retaining only variants reported in *≥* 3 independent studies and *≥* 3 bacterial species. Following this curation, we obtained a final panel of eight mutations covering five positions in the *rpsL* gene: 43, 86, 88, 91, and 92 (Figure 1). Among these, 43R and 88R were the most frequently reported, detected in 27 and 18 species, respectively. Position 43 with three variants, 43R, 43N, and 43T, was the most mutation-prone site, collectively appearing in a total of 49 species (Figure 1A). With variations found in both lab and natural isolates, *M. tuberculosis* and *E. coli* exhibited the greatest mutational diversity. However, mutations in *Thermus thermophilus* were exclusively laboratory-derived, while those in *Klebsiella pneumoniae* were reported only in clinical isolates (Figure 1B).

**Figure 1.**
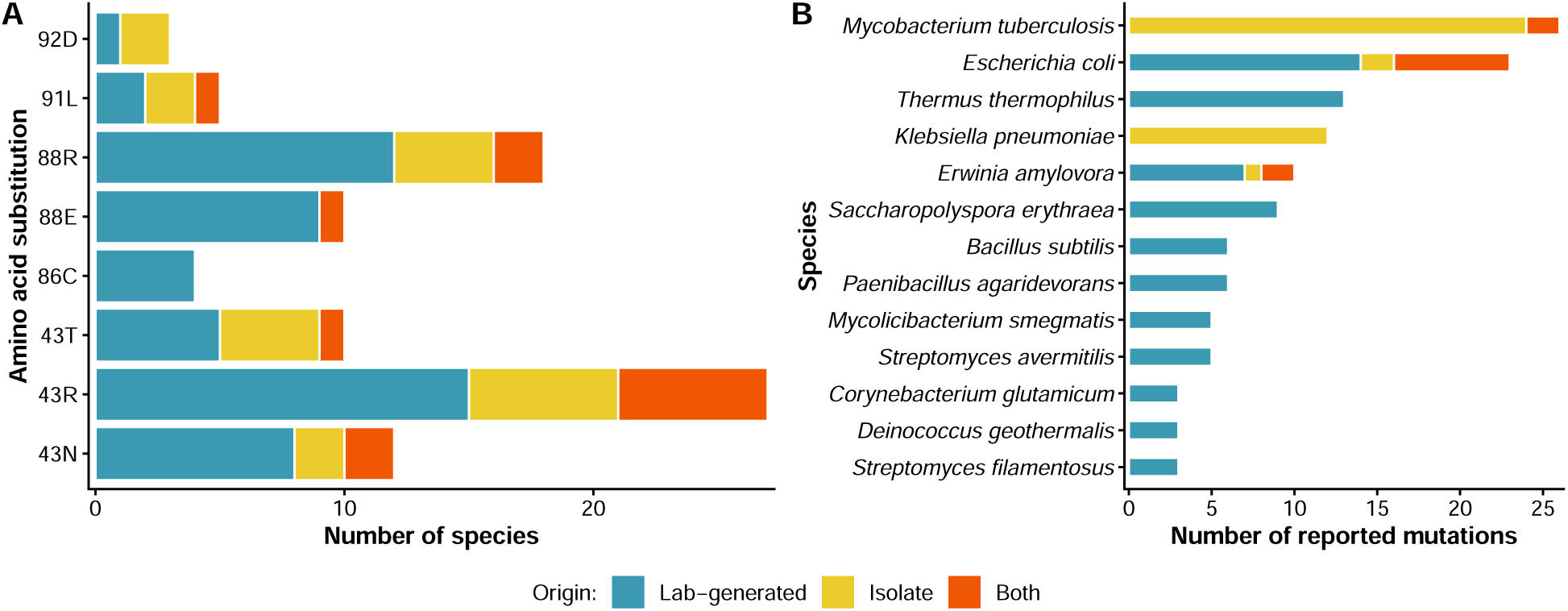
STR resistance mutations identified in *rpsL* from published studies. A) Number of species in which each amino acid substitution was reported. Only substitutions observed in at least 3 species are shown. B) Number of reported mutations per species, with only species having *≥* 3 substitutions included. Colors indicate mutation origin: laboratory-generated (blue), clinical/environmental isolates (yellow), or both (orange). **Alt text:** Two-panel bar chart of *rpsL* streptomycin resistance mutations from published studies. Panel A shows species counts per amino acid substitution; Panel B shows reported mutations per species, color-coded by mutation origin.

Using the same approach for *rrs*, we compiled a database of STR resistance–conferring mutations from 32 studies, which together reported a total of 111 *rrs* mutations in five species. By merging homologous residues, we identified 49 unique nucleotide substitutions at 41 positions. The majority of the reported mutations were observed in *M. tuberculosis* and were predominantly derived from isolate-based studies (Figure S1). Since fewer studies reported *rrs*-associated mutations, a less stringent filter was applied, resulting in four high- confidence mutations: 523C, 885A, 912G and 913G. All selected mutations were supported by evidence from both clinical isolates and laboratory-generated experiments.

To visualize the structural relevance of the selected mutations, we mapped the five *rpsL* and four *rrs* sites from our screening panel onto a ribosome structure of *E. coli*, focusing on both the ribosomal protein S12 (*rpsL*) and 16S rRNA (*rrs*) (Figure S2). This structure is derived from the conserved antibiotic-ribosome interactions of *E. coli* (Paternoga et al. 2023) in the RCSB Protein Data Bank (PDB) (Berman 2000). The figure illustrates the close spatial proximity between the STR molecule and the key sites identified in this study.

### Phylogenetic distribution of predicted intrinsic and evolvable STR resistance across bacteria taxa

#### Ribosomal protein S12 (*rpsL*)

We extracted 20 808 *rpsL* sequences from 20 806 annotated bacterial reference or representative genomes from the NCBI database. After applying quality control filters, a total of 15 740 *rpsL* sequences of 15 721 species were retained (Figure S3). Only 15 genomes contained more than one *rpsL* sequence, with a maximum of two copies per genome.

The filtered *rpsL* sequences were then screened for the presence and evolvability of eight mutations from our panel of reported resistance to STR. In total, 1190 species (7.57% of the analyzed dataset) are predicted to exhibit intrinsic resistance to streptomycin on the basis of harboring at least one of these eight mutations. These species were then assigned to the GTDB (Parks et al. 2022)-based phylogenetic tree comprising 12 164 species. Several distinct phylogenetic clusters of predicted intrinsic resistance are evident (permutation test, *p <* 0.001), including the orders Sphingomonadales and Rickettsiales, the family Devosiaceae (all within the Alphaproteobacteria), and the classes Coriobacteriia and Planctomycetia (Figure 2). In contrast, intrinsically resistant species are largely absent from several major clades, including the Bacteroidia and Gammaproteobacteria classes (Figure 3A).

**Figure 2.**
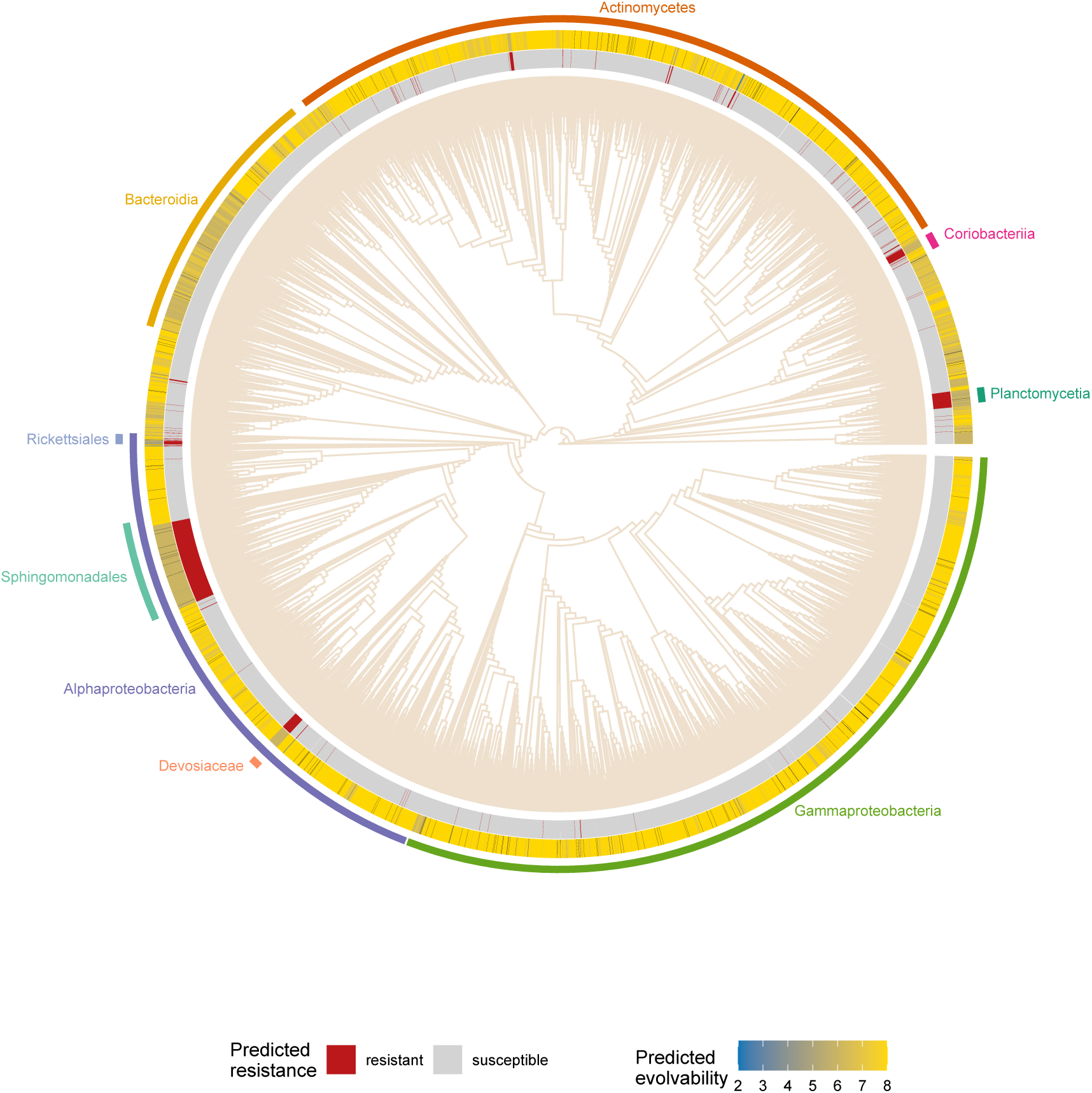
Phylogenetic distribution of predicted intrinsic resistance and evolvability of streptomycin resistance based on *rpsL*. The tree represents a subset of the full bacterial phylogeny from the Genome Taxonomy Database (Parks et al. 2022). Red segments in the inner circle indicate species predicted to be intrinsically resistant, defined by the presence of at least one *rpsL* resistance-associated amino acid substitution from our curated panel. The outer ring shows the predicted evolvability, defined as the number of resistance mutations accessible via single-nucleotide changes. **Alt text:** Circular phylogenetic tree showing predicted intrinsic resistance and evolvability based on *rpsL* across bacteria, with an inner ring marking intrinsic resistant clades and an outer ring showing evolvability scores across bacterial taxa.

**Figure 3.**
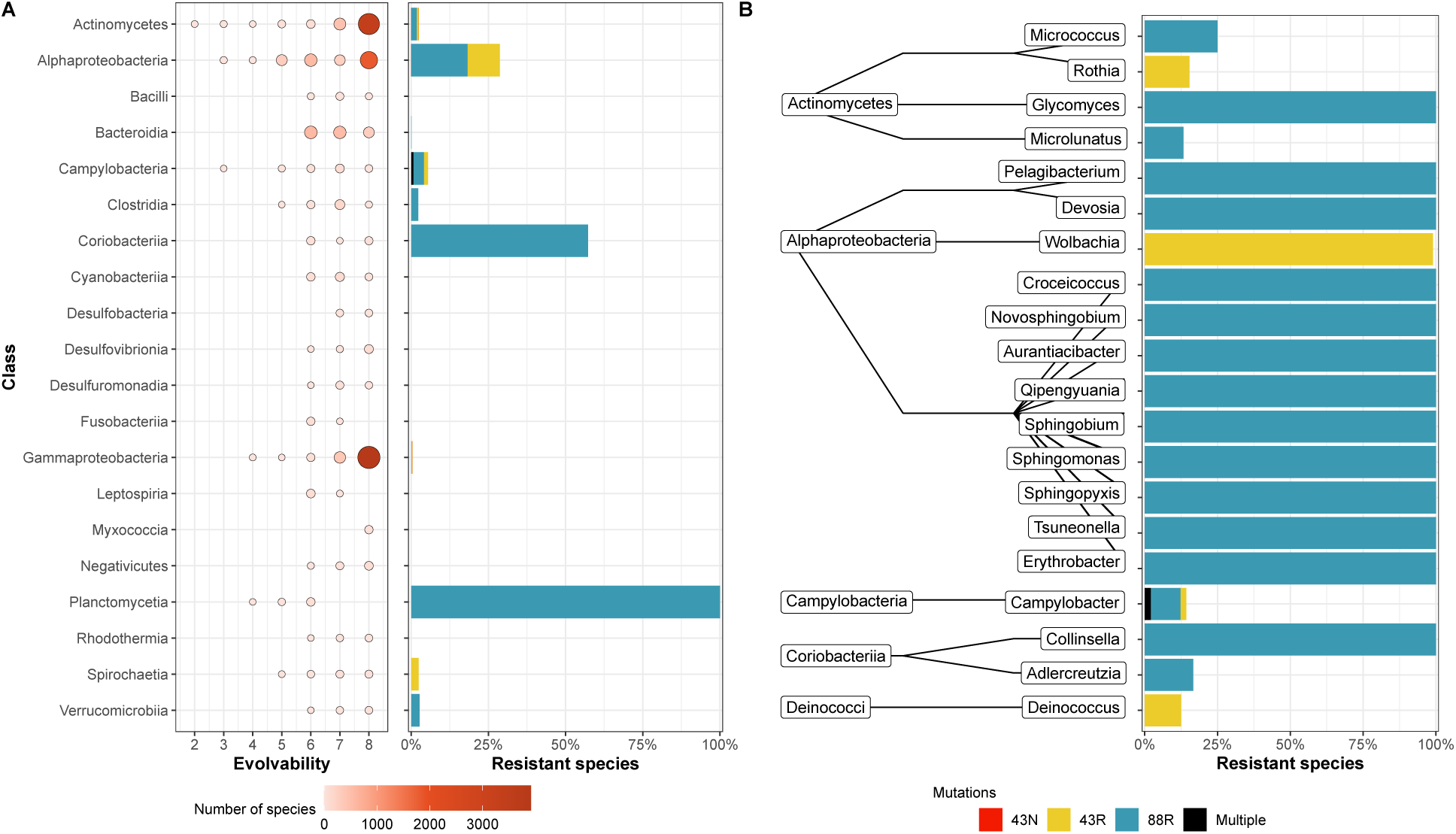
Predicted evolvability and intrinsic resistance across taxonomic levels based on *rpsL*. (A) Evolvability of species across the twenty most speciose bacterial classes is shown as bubble plots, where the x-axis indicates the number of possible resistance-conferring mutations. Bubble size and color intensity correspond to the number of species within each category. The percentage of species predicted to be intrinsically resistant across the same class is shown as a bar plot, with color-coding indicating specific resistance mutations (43N, 43R, 88R, or multiple). (B) Percentage of intrinsic resistance among genera with the highest resistance rates, grouped by class. Only genera containing *≥*10 species were included (except for *Wolbachia*, for which we analysed 293 genomes from different hosts but which has only one officially recognized species). **Alt text:** Two-panel figure showing streptomycin resistance predictions based on *rpsL*. Panel A: bubble plots of evolvability (number of accessible resistance mutations, 2–8) across 20 bacterial classes, paired with bar graphs showing percentage of intrinsically resistant species of the same class. Panel B: bar chart of intrinsic resistance percentages in top genera grouped by class, with multiple genera in Alphaproteobacteria showing the highest proportion of resistant species.

Evolvability, in the context of this study, refers to the number of STR resistance mutations at the amino acid level that can arise from a single nucleotide substitution. This resistance capacity to evolve varies across bacterial species, with values ranging from 2 to 8 possible mutations (Figure 2). However, evolvability is largely constrained within a narrow range, with 95% of species falling between 6 and 8 accessible mutations. A similar pattern is observed across the 20 most represented bacterial classes (Figure 3A). Statistical analysis indicates a strong phylogenetic signal in evolvability (Pagel’s *λ* = 0.991, *p <* 0.001; Blomberg’s *K* = 0.372, *p* = 0.001).

Based on our *rpsL* screening, approximately 70% of intrinsically resistant species carry the 88R mutation. Only seven species with mutations other than 88R or 43R are identified, including substitutions 43N and 43T or combinations of mutations. Planctomycetia exhibits the highest proportion of predicted resistant species (100%), with all 70 species carrying the conserved 88R mutation. Alphaproteobacteria contain the largest number of intrinsically resistant species overall (848 species), driven by three key clades Sphingomonadales (472), Rickettsiales (293), and Devosiaceae (59). Within the Rickettsiales, 290 of the 293 screened *Wolbachia* genomes are predicted to be intrinsically resistant. Notably, species from the genus *Rickettsia* are not shown in the Rickettsiales phylogeny, as the majority were excluded due to high divergence in the *rpsL* sequences relative to the reference (Figure S4). Additional analysis indicates that none of the excluded Rickettsia genomes carries resistance-associated mutations. Other classes comprising many species predicted to be intrinsically resistant to STR include Actinomycetes (96 species) and Coriobacteriia (55 species). Figure 3B presents the phylogenetic distribution of intrinsic resistance across dominant bacterial classes, also highlighting genera with the highest proportion of resistant species.

#### 16S rRNA (*rrs*)

Compared to *rpsL*, substantially fewer high-confidence *rrs* mutations have been reported. This limited mutation panel restricts our ability to draw phylogenetic inferences from *rrs*. Nevertheless, we performed a comprehensive screen of all *rrs* sequences for four curated resistance mutations to identify intrinsically resistant species. We find that intrinsic resistance inferred from *rrs* screening is extremely rare. Only eight species (approximately 0.04%), spanning six genera and predominantly within the Gammaproteobacteria class, are predicted to carry these resistant variants (see Supplementary Material for details).

#### Mutational landscape of STR resistance evolvability

To investigate how the evolvability of individual STR resistance mutations varies across bacterial species, we examined the mutational possibility for eight key amino acid substitutions in *rpsL* (Figure 4). Mutations 43R, 43T, 88R, and 91L can easily arise in almost all bacterial species by a single-nucleotide substitution. In contrast, mutations 92D, 88E, 86C, and 43N are only accessible to a subset of species. 86C stands out as the most constrained, with the highest number of species predicted to be unable to evolve this mutation. The evolvability of these substitutions also varies across bacterial phyla and classes (Figure S5). In addition, figure 4 highlights intrinsically resistant species, including 830 carrying the 88R variant and 359 carrying the 43R variant. Meanwhile, 360 species are predicted to be unable to evolve the 43N mutation through a single nucleotide substitution. These observations raise the question whether different mutations that occur at the same codon position are mutually exclusive, so that the presence of one variant constrains the accessibility of alternative resistance mutations.

**Figure 4.**
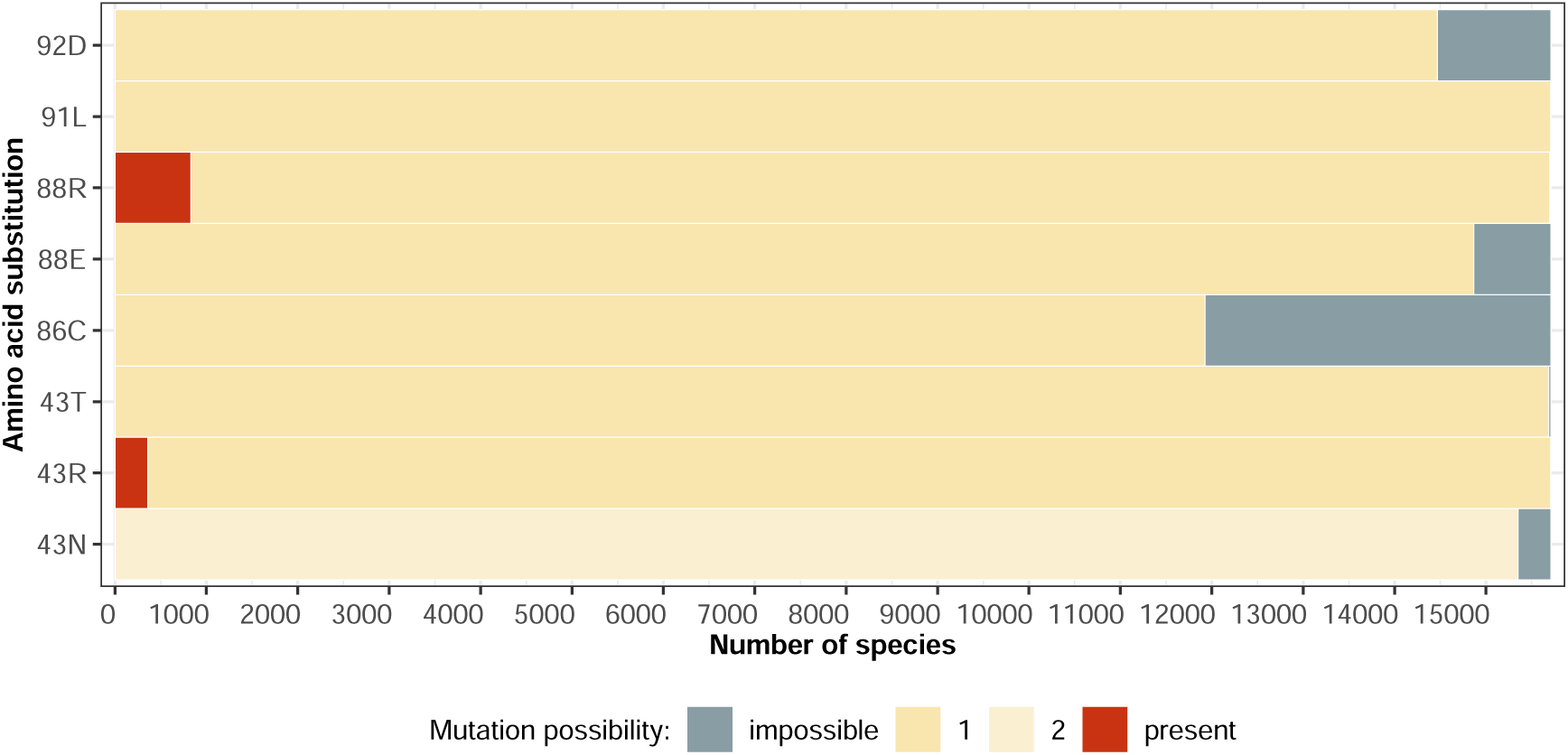
Mutational landscape of intrinsic and evolvable STR resistance through *rpsL* mutations. The plot shows the distribution of STR resistance mutations across amino acid residues targeted in our screened panel. For each residue, species are categorized based on whether the resistance mutation is: (i) already present (red), (ii) potentially evolvable via a single nucleotide substitution (yellow), or (iii) impossible to evolve through a single nucleotide change (grey). The intensity of yellow indicates evolvability, depending on whether the mutation can arise from one or two distinct point mutations at the corresponding codon. Species with multiple *rpsL* sequences were excluded from the plot. **Alt text:** Stacked bar chart showing mutational landscape of *rpsL* streptomycin resistance across approximately 15 000 species, categorizing each of eight mutations as present, evolvable via one or two nucleotide substitutions, or inaccessible.

To further elucidate the observed variation in evolvability, we examined the underlying mutational codon networks. These networks are graphs representing all possible codon states at a given amino acid position, where edges connect codons that are interconvertible through single nucleotide substitutions. Figure 5A shows an example for position 86. At this site, a single amino acid residue (cysteine) has been reported to confer STR resistance (86C). Although 99% of species in our data set encode arginine (R) at this position, the use of codons varies: the most frequent codons are CGT and CGC (at 60% and 16%, respectively), among other arginine codons. Only these two codons can mutate toward cysteine codons (TGT and TGC) in a single mutation step, which explains why approximately 24% of species cannot evolve resistance via a single step mutation at this position.

**Figure 5.**
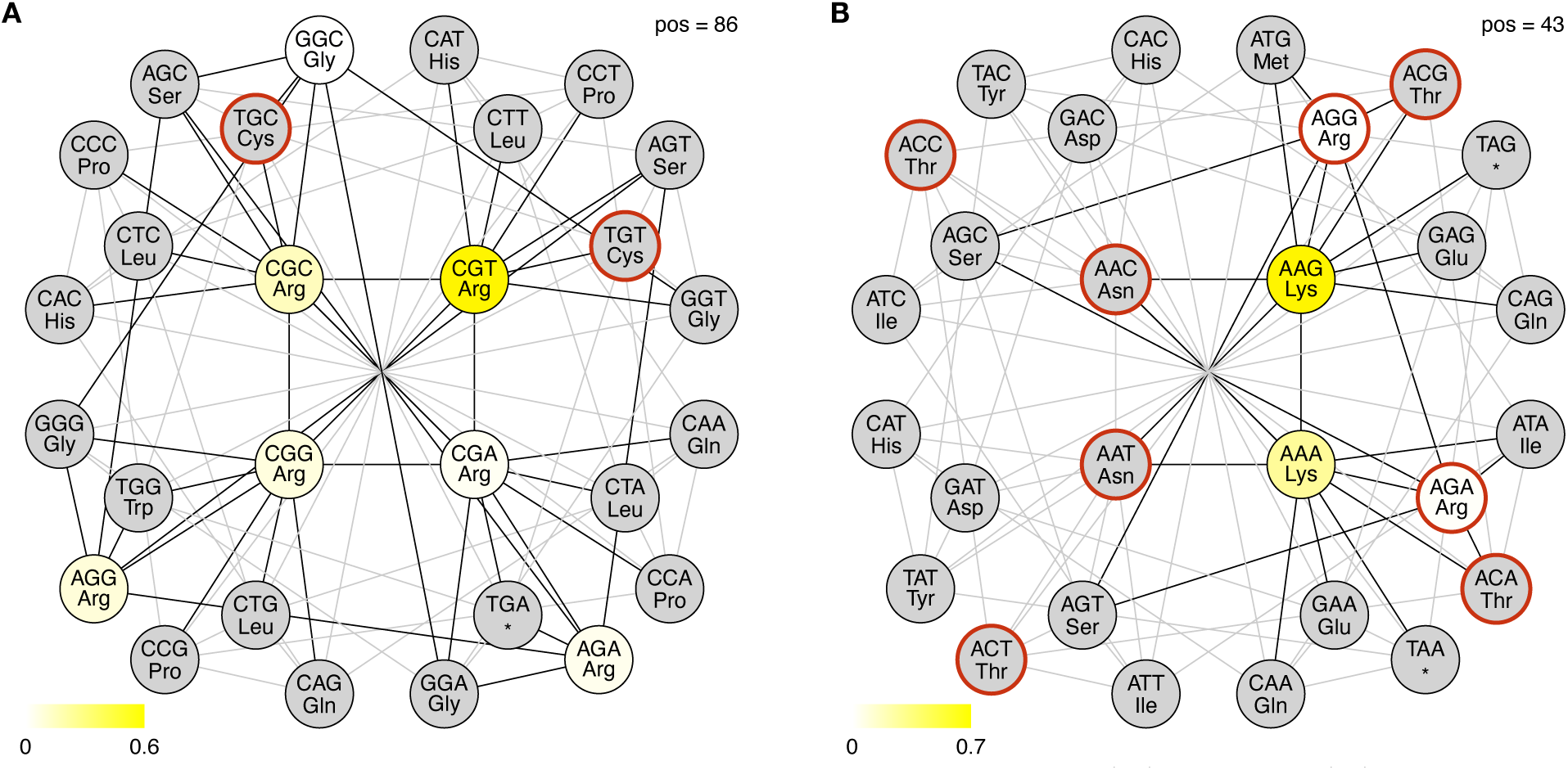
Codon network for *rpsL* amino acid positions 86 (A) and 43 (B), illustrating mutational accessibility. Each circle represents a codon, with edges connecting codons that differ by a single nucleotide substitution. The yellow color gradient indicates the proportion of species encoding each codon, while grey indicates codons not found in any species. Red circles mark codons associated with STR resistance. Edges connecting observed codons are shown in black, while edges between unobserved codons are shown in grey. **Alt text:** Codon network diagrams for *rpsL* amino acid positions 86 and 43, subfigures A and B, illustrating single-nucleotide mutational pathways between codons, the frequency of each codon across species, and which codons are associated with streptomycin resistance.

A more complex scenario occurs at position 43, for which three resistance-associated amino acid substitutions—arginine (R), asparagine (N), and threonine (T)—have been reported (Figure 5B). At this position, 98% of the species encode lysine (K), represented by two codons AAG (70%) and AAA (28%). Each lysine codon can mutate to asparagine codons (AAC and AAT) via two distinct single nucleotide changes, while only one mutation pathway exists to convert to arginine (AGG or AGA) or threonine (ACG or ACA). There are no connecting edges between the arginine (R) and asparagine (N) codons in the network. Therefore, a species that has the substitution 43R cannot evolve 43N through asingle-step mutation. In contrast, arginine can mutate to threonine through a single-nucleotide change, meaning that this constraint does not apply to 43T. A similar constraint is observed at position 88. The network for position 88 (Figure S6) shows that there is no pathway between arginine (R) and glutamic acid (E). Thus, species carrying the 88R mutation cannot evolve to 88E by a single nucleotide change.

## Discussion

Our results indicate that the overall evolvability of STR resistance through target modification mutations varies very little across the bacterial tree of life. Most species have evolutionary access to between six and eight out of the eight *rpsL* mutations in our panel, indicating that there are only limited genetic constraints on STR resistance evolution. Nevertheless, evolvability varies across individual mutations, and this variation can result in very different phenotypic outcomes. Mutations that confer antibiotic resistance affect several essential processes and often impose fitness costs; however, these costs differ widely among individual variants. For example, STR resistance mutations K43R and K88R impose low or no fitness cost across multiple species, including *Mycobacterium smegmatis*, *Salmonella enterica* serovar Typhimurium, *Erwinia amylovora*, and *E. coli* (Björkman et al. 1998; Sander et al. 2002; Escursell et al. 2021; Hinz et al. 2024). In contrast, K43N, K43T and K88E mutations significantly reduce fitness in these same species (Björkman et al. 2000; Sander et al. 2002; Trindade et al. 2009; Escursell et al. 2021). The apparent clustering of mutations into high-and low-cost groups across several species contrasts with observations for other antibiotics where fitness cross are less predictable across different genetic backgrounds (Vogwill et al. 2016; Card et al. 2021; Hinz et al. 2024; Williams et al. 2025). The mutation-specific fitness differences may also explain the uneven distribution of intrinsic resistance across bacterial taxa. Approximately 7.6% of all species carry at least one *rpsL* resistance mutation, yet nearly all of these species harbor one of only two variants (43R or 88R). Given that both K43R and K88R have been reported to impose little or no fitness cost in several bacterial species, their high prevalence may reflect enhanced long-term persistence and broader evolutionary success relative to more costly alternatives.

Beyond their individual effects, *rpsL* mutations can vary in how they interact epistatically with other resistance mutations. For example, positive epistasis has been detected between K43R and D87G in *gyrA* (conferring resistance to nalidixic acid), while combinations of *rpsL* K43T/K43N with *gyrA* D87G show negative epistasis (Trindade et al. 2009). Similarly, both positive and negative epistatic interactions have been reported between *rpsL* mutations and various rifampicin resistance mutations in *rpoB*. While no significant epistatic interaction is observed between K43R and the H526Y *rpoB* variant, the K43N substitution shows positive epistasis with the same variant (Trindade et al. 2009). Conversely, a strong negative epistatic interaction has been documented between K43T and H526Y (Trindade et al. 2009; Moura De Sousa et al. 2017).

Intrinsic resistance to STR is predicted in several clades, including Sphingomonadales, Devosiaceae, Rickettsiales, Coriobacteriia, and Planctomycetia. Three of these (Sphingomonadales, Devosiaceae, and Rickettsiales) belong to the Alphaproteobacteria class, of which the Sphingomonadales are the largest. In accordance with this prediction, there is experimental evidence demonstrating that arginine at position 88 (R88) in *rpsL* confers streptomycin resistance in *Sphingomonas* sp. strain Fr1, as the substitution of this residue with lysine (R88K) abolishes resistance (Kaczmarczyk et al. 2012). Our analysis also detects the 88R mutation in all examined species in the Planctomycetia class. Several members of the Planctomycetia phylum are commonly isolated using medium supplemented with ampicillin and streptomycin, to which many planctomycetal species are known to be resistant (Lage and Bondoso 2011; Wiegand et al. 2019). Within the Rickettsiales, we found that in addition to *Wolbachia* genomes, *Ehrlichia ruminantium* also carries an STR mutation (43R) (Figure S4). This pattern is consistent with previous research that identified the variant K43R in *Wolbachia* and *E. ruminantium*, but not in related genera such as *Anaplasma* and *Neorickettsia* (Fallon et al. 2013). The evolutionary drivers behind this variant in these lineages, particularly in *Wolbachia*, remain elusive. As an obligate endosymbiont of arthropods and nematodes, *Wolbachia* replicates only within eukaryotic host cells where exposure to aminoglycosides such as streptomycin is expected to be limited. However, aminoglycoside-producing *Streptomyces*, a genus of more than 500 species, are abundant in soil and decaying vegetation. The degree to which insects and their associated microbiota encounter these antibiotics in natural environments remains poorly characterised (Fallon et al. 2013).

Intrinsic resistance based on 16S rRNA (*rrs*) variants appears to be extremely rare. Among 45 000 *rrs* sequences from approximately 19 000 surveyed species, only 8 species (0.04% of the data set) carry mutations included our panel. These species are distributed in six genera and predominantly within the Gammaproteobacteria (see Supplementary Material for details). These *rrs* mutations are uncommon since 16S rRNA forms a highly conserved core of the ribosomal decoding center and mutations that affect this essential machinery frequently result in cell death (Garneau-Tsodikova and Labby 2016). The prediction of intrinsic resistance based on *rrs* is further complicated by the multiplicity of the rRNA gene in many genomes. Only one species harbors a single *rrs* copy predicted to be resistant, whereas the remaining seven species possess more than two copies. In such contexts, wildtype alleles may mask the phenotypic effect of a resistant copy, limiting both selection for resistance via these variants and the reliability of *rrs*-based predictions of intrinsic resistance.

It is important to stress that although the presence of resistance mutations in *rpsL* and (to a lesser degree) *rrs* enables predictions of STR resistance, absence of these mutations does not predict susceptibility to STR. This is because STR resistance can be caused by a variety of resistance mechanisms other than target modification. *Streptomyces griseus*, the bacterium from which STR was first derived, is a case in point. Even though this bacterium produces STR it does not harbour any of the *rpsL* or *rrs* resistance mutations in our panel. Instead, resistance is achieved through the expression of two streptomycin phosphotransferases (SPH(6) and SPH(3”)) that inactivate the drug (Miller and Walker 1969; Nimi et al. 1971; Cundliffe 1989). Similarly, the *Rv2004c* gene, a member of the DosR regulon, confers low-level resistance to STR through phosphotransferase activity in *M. tuberculosis* (Doddam et al. 2019). Another notable example involves mutations in *gidB*, which encodes a 16S rRNA methyltransferase. Mutations in *gidB* confer low-level STR-resistance (Okamoto et al. 2007), and a point mutation in *gidB* has been reported to confer intrinsic STR resistance in *Bacillus velezensis* (Na et al. 2025). However, the association of mutations in *gidB* with STR resistance needs to be further investigated, as several *gidB* mutations have been found in both susceptible and resistant isolates (Spies et al. 2011; Feuerriegel et al. 2012).

STR is an aminoglycoside antibiotic, a class of compounds that comprises an amino alcohol ring and amino sugar molecules linked by a ligand sugar bond. Numerous aminoglycosides used clinically today include natural aminoglycosides (e.g., STR, kanamycin, and gentamicin), as well as semi-synthetic derivatives (e.g., amikacin, etimicin, and netilmicin). Intuitively, one might expect that at least some of the resistance mutations in our panel also confer cross-resistance to other aminoglycosides. However, this is not generally the case. Streptomycin is an atypical aminoglycoside both in terms of structure (containing a streptidine instead of a deoxystreptamine ring) and binding (binding to a different position of the 16S rRNA where it also interacts with the S12 protein). As a result, mutations in *rpsL* do not confer resistance to other aminoglycosides. Some limited degree of overlap between mutations in *rrs* conferring resistance to different aminoglycosides appears to exist. For example, *rrs* A1401G has been reported in isolates of *M. tuberculosis* resistant to both STR and amikacin (Islam et al. 2020), and A1408G has been reported to confer both STR and kanamycin resistance in *Thermus thermophilus* (Gregory et al. 2005). However, the same mutation only confers kanamycin and gentamicin but not STR resistance in *Borrelia burgdorferi* (Criswell et al. 2006). In general, however, there is limited scope for *rrs*-mediated cross-resistance because target-site mutations the 16S rRNA are a rare mechanism of resistance to aminoglycosides in fast-growing bacteria (Garneau-Tsodikova and Labby 2016).

Mirroring the results of a recent study of rifampicin resistance by Bolourchi *et al*. (Bolourchi et al. 2025), our results indicate that STR resistance is highly evolvable across the bacterial tree of life. Both studies suggest that genetic constraints on resistance evolution are generally limited. In the rifampicin study, most species had evolutionary access to a large set of one-step *rpoB* resistance mutations (typically 35–47 of 57 mutations screened), whereas in our STR analysis, most species can access 6–8 of the eight characterised *rpsL* mutations. In terms of intrinsic resistance, both antibiotics exhibit clear phylogenetic clustering, but with enrichment in different clades. Approximately 8% of the species were intrinsically resistant to rifampicin; they were often concentrated in Spirochaetia, Mollicutes, and many members of the Actinomycetes class. By contrast, we predict 7.6% of species carry intrinsic STR-resistance mutations, with distinct hotspots in Alphaproteobacteria (e.g., Sphingomonadales, Rickettsiales) and the class Planctomycetia. Beyond taxonomic differences, the genetic architecture of intrinsic resistance also differs between drugs: while STR resistance is more restricted and heavily dominated by a small number of low-fitness-cost mutations (43R and 88R), rifampicin resistance mutations in *rpoB* are more broadly distributed across mutation types.

Our study has several limitations. First, to build a panel of resistance mutations for STR, it is necessary to strike a balance between comprehensiveness (capturing as many reported mutations as possible) and robustness (ensuring that the included mutations are reliably associated with resistance to STR). To improve robustness, we filter mutations based on both the number of supporting studies and the diversity of species in which they were reported. This limitation is particularly evident for *rrs*, for which few studies—most of them in *M. tuberculosis*—report resistance-associated mutations, resulting in a limited set of candidates with high-confidence. However, this conservative approach ensures that our predictions are grounded in well-validated determinants, thus minimizing the risk of false positives. Consequently, future efforts in the curation of data and the experimental validation of resistance markers in a wider range of bacterial lineages will be essential to expand this catalog. Second, our analysis used a single representative genome per species, which may not fully capture intraspecific variation. As a result, certain species predicted as susceptible may, in fact, harbor resistant lineages that were not represented in our dataset. Future investigations employing pangenomic datasets will be necessary to characterize the full extent of this diversity. Additionally, a focus on highly divergent or specialized groups, such as *Rickettsia*, is warranted. Their extreme sequence divergence suggests that, while reference-based screening is effective for the majority bacterial clades, these unique lineages may require clade-specific models to accurately predict resistance.

In conclusion, our computational framework provides a global view of STR resistance across the bacterial tree of life by integrating known resistance-associated variants with codon-level mutational accessibility. We predict significant levels of intrinsic resistance in Alphaproteobacteria, Coriobacteriia, and Planctomycetia. Furthermore, our prediction of evolvability indicates that most bacterial species can easily access most of the high-confidence resistance mutations in our panel, suggesting limited genetic constraints on the future emergence of STR resistance. Although experimental validation remains necessary, by identifying existing natural reservoirs and quantifying the future evolutionary potential of resistance, this work establishes a predictive framework to explore the emergence and evolution of STR resistance across bacteria.

## Acknowledgements

We thank Negin Bolourchi for helpful discussions and guidance on adopting her bioinformatics pipeline.

