## Supplementary Material for "Mutations associated with streptomycin resistance predicted to be highly prevalent and evolvable across bacteria"

### S1. Supplementary Figures

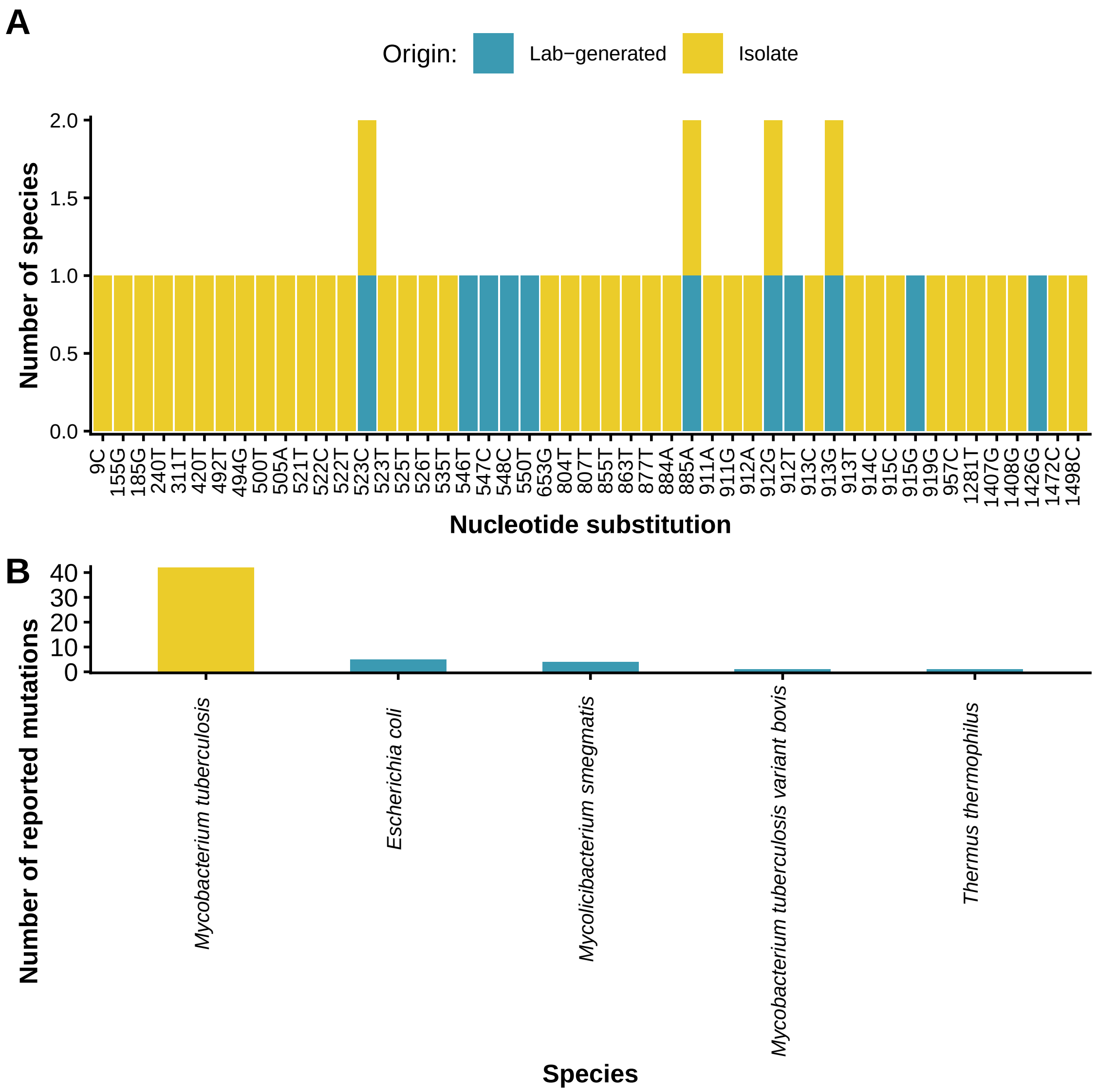


**Figure S1.** STR resistance mutations identified in *rrs* from published studies. A) Number of species that nucleotide mutations were reported in. B) Number of reported mutations per species. Colors indicate mutation origin: laboratory-generated (blue), clinical/environmental isolates (yellow).

**Alt text:** Bar charts of *rrs* streptomycin resistance mutations from published studies, subfigures A and B showing species counts per nucleotide substitution and reported mutations per species, dominated by *Mycobacterium tuberculosis*.

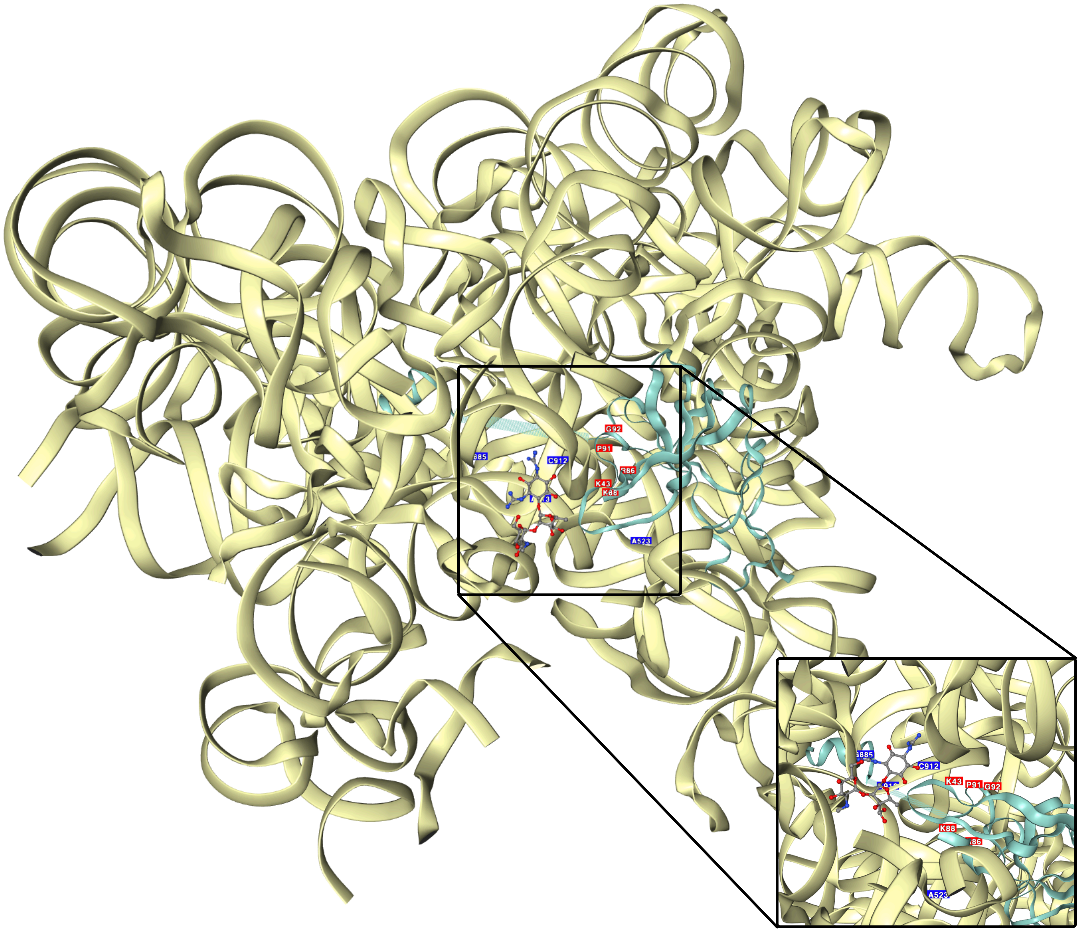


**Figure S2.** Structural visualization of STR resistance mechanisms from the RCSB Protein Data Bank (PDB) (Berman 2000), highlighting the two key gene products: ribosomal protein S12 (*rpsL*, green) and 16S rRNA (*rrs*, yellow). STR, shown in a stick-and-ball representation, binds to the 16S rRNA. Mutations in *rpsL* and *rrs* alter the binding site, leading to resistance. Resistance-associated substitution sites in *rpsL* are shown in red, those in *rrs* are also shown in blue, and the STR-binding region is outlined by a black square. The inset provides a close-up view of mutations located and STR. An interactive version of this visualization is available in the GitHub repository under `plots/rpsL_rrs_structure.html`.

**Alt text:** Three-dimensional structure of the *E. coli* ribosome showing streptomycin binding to ribosomal protein S12 and 16S rRNA, with resistance-associated sites highlighted and an inset close-up of the binding interface.

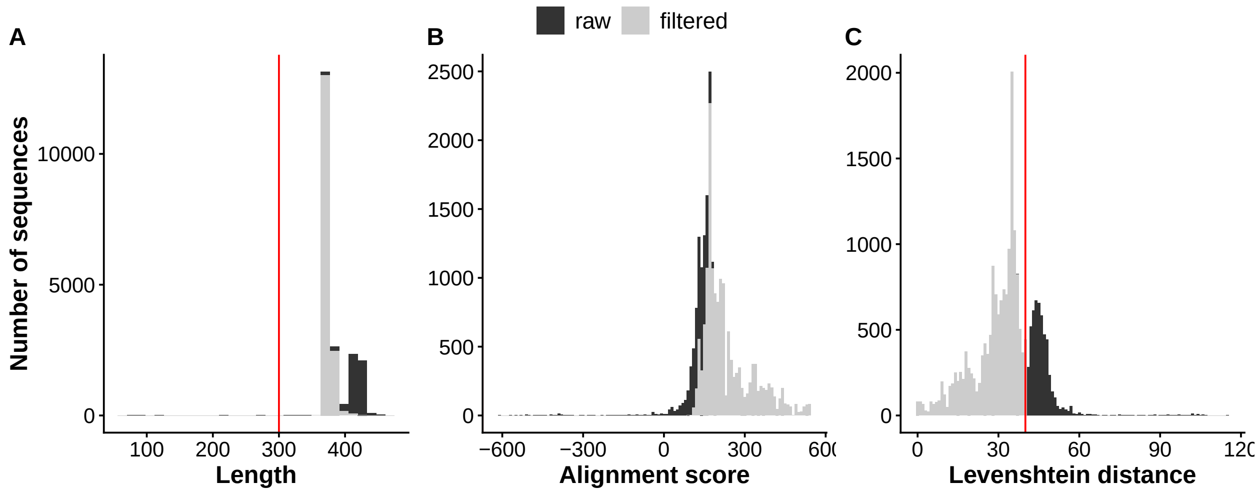


**Figure S3.** Properties of *rpsL* sequences extracted from genomes downloaded from NCBI. The plots show the distribution of key sequence properties before and after (grey) filtering. A) Distribution of sequence lengths, with a vertical red line indicating the minimum length threshold of 300 bp. B) Alignment scores of each sequence against the *E. coli rpsL* reference sequence. C) Levenshtein distance of the entire sequence relative to the *E. coli* reference, with a red line indicating the maximum threshold of 40 used for filtering.

**Alt text:** Histograms of *rpsL* sequence properties before and after quality filtering, subfigures A, B and C showing distributions of sequence length, alignment score, and Levenshtein distance relative to the *E. coli* reference, with filtering thresholds indicated.

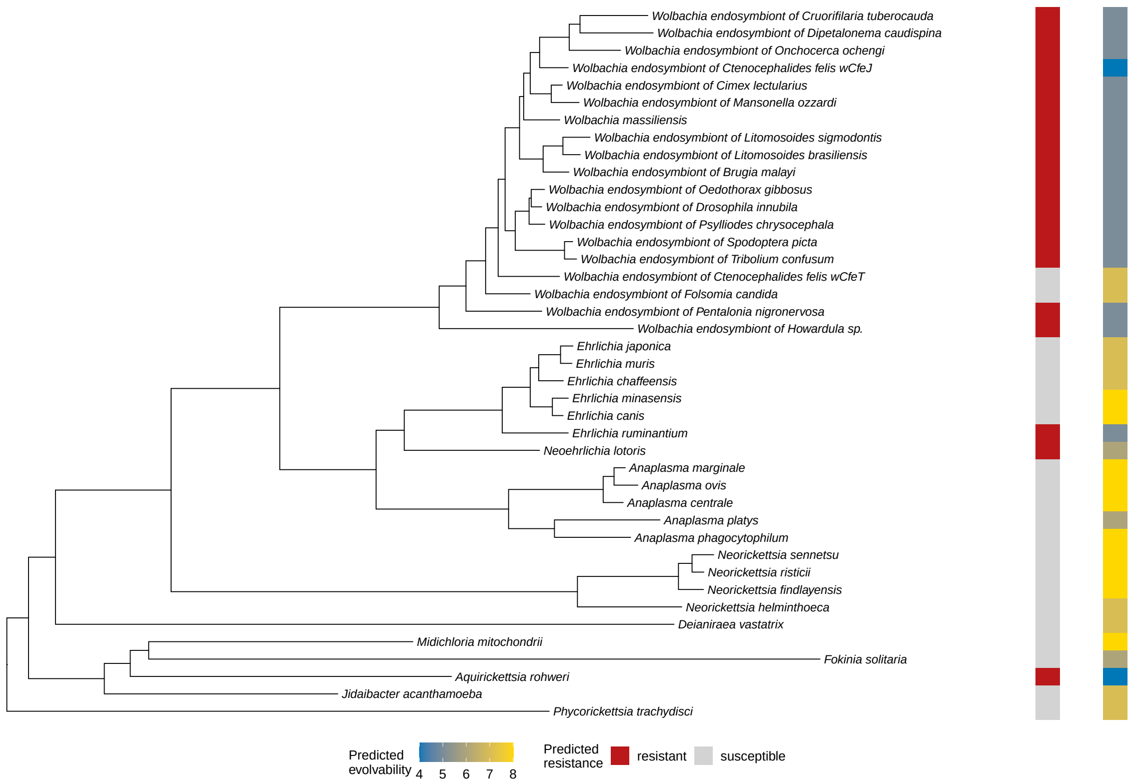


**Figure S4.** Phylogenetic tree of the order Rickettsiales. Predicted intrinsic resistance is indicated in the same way as in Figure 2. No *Rickettsia* species are shown, as most were excluded due to high Levenshtein distances from the reference sequence.

**Alt text:** Phylogenetic tree of the order Rickettsiales showing predicted intrinsic streptomycin resistance and evolvability, with no *Rickettsia* species shown due to high sequence divergence.

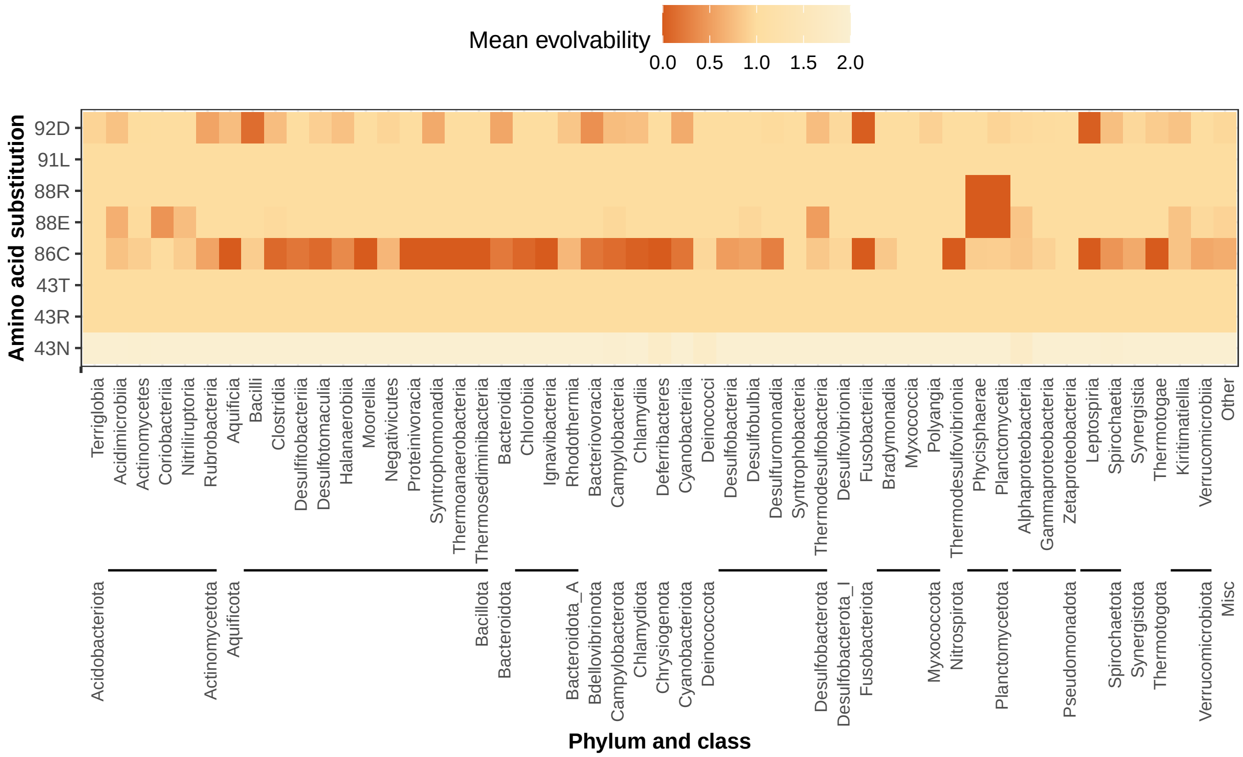


**Figure S5.** Mean evolvability of STR resistance mutations across bacterial classes based on *rpsL*. Evolvability is defined as the mean number of nucleotide substitutions per species that can give rise to a specific amino acid substitution conferring resistance. Bacterial classes are grouped by phylum. Species already harboring resistance-associated mutations were excluded from the figure.

**Alt text:** Heatmap of mean *rpsL* evolvability for eight resistance mutations across bacterial classes grouped by phylum, showing variation in mutational accessibility across the bacterial tree of life.

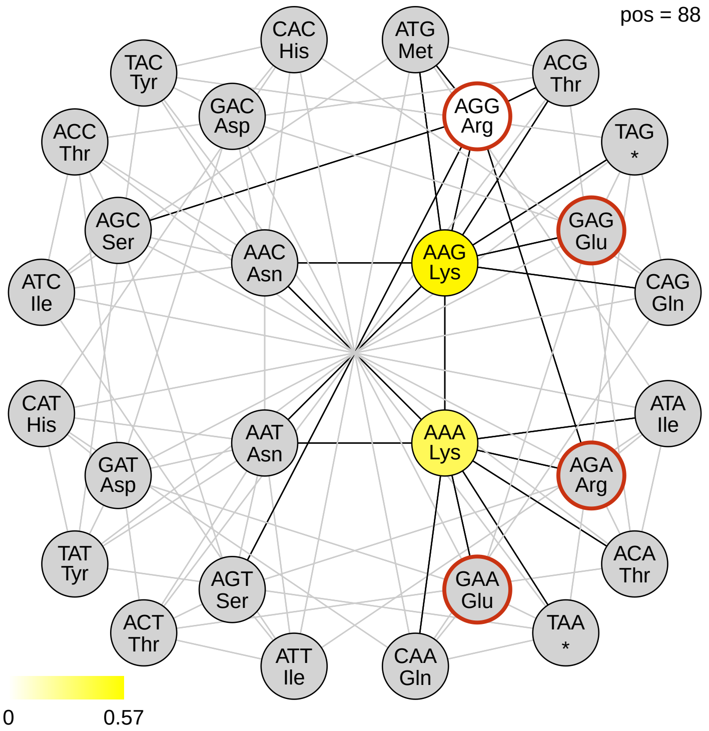


**Figure S6.** Codon network at *rpsL* amino acid position 88. All visual elements and color coding are consistent with Figure 5.
**Alt text:** Codon network diagram for *rpsL* amino acid position 88, illustrating single-nucleotide mutational pathways between codons, the frequency of each codon across species, and which codons are associated with streptomycin resistance.

### S2. Supplementary Results

A total of 46 306 *rrs* sequences were extracted from the NCBI database. Following quality control, 45 057 sequences representing 18 926 species were retained (Figure S7). The filtered *rrs* sequences were then screened for the presence of four resistance mutations. Only 8 species (0.04% of the species in our dataset) were found to carry at least one *rrs* resistance mutation and none of these species harbour *rpsL* resistance mutations. These species are distributed in six genera, but are predominantly concentrated within the Gammaproteobacteria class (Table S1). Unlike *rpsL*, *rrs* evolvability was not assessed, as *rrs* is not codon-based and every resistance mutation is evolvable in a single-step mutation.

Among species predicted to be intrinsically resistant based on *rrs* mutations, only a minority of *rrs* gene copies carried the resistant variant, despite many of these species harboring multiple *rrs* operons (Table S1). *Zobellia laminariae* has previously been reported to exhibit resistance to STR (Nedashkovskaya et al. 2004). In addition, some *Pectobacterium* strains have been described as naturally resistant to STR (Vu et al. 2022). All *Vibrio* species isolated from the Andaman Sea were reported to have resistance to STR; however, *Vibrio aquimaris* was not included in that study, limiting direct comparison with our predictions. Unlike other intrinsically resistant species that harbor multiple *rrs* operons, *Spiroplasma taiwanense* contains only a single *rrs* copy, which carries resistance mutations. While direct phenotypic evidence for STR resistance in *S. taiwanense* is absent, reduced STR susceptibility has been reported in other *Spiroplasma* species, such as *Spiroplasma citri* (Liao 1981).

In contrast to *rpsL*, a substantially larger number of genomes (7456) in our data set contain multiple gene copies, ranging from one to 37 per genome. Variation in *rrs* copy number has been widely documented (Stoddard et al. 2015; Větrovský and Baldrian 2013). rRNA copy number variation is often associated with the strategy of bacterial life. Species with a higher number of copies are often able to respond quickly to favorable growth conditions, whereas those with fewer copies are typically associated with oligotrophic lifestyles (Roller et al. 2016). Although the distribution of the *rrs* copy number is highly skewed, with most genomes carrying a single gene copy, a substantial fraction of genomes have multiple operons (Figure S8). This multicopy architecture introduces within-genome heterogeneity as resistance mutations present in one operon may be buffered by co-existing wild-type copies, limiting their phenotypic effect. As a result, the presence of a resistant *rrs* allele alone may not be sufficient to reliably predict phenotypic STR resistance. Consequently, in contrast to single-copy protein-coding targets such as *rpsL*, where resistance mutations can have immediate and dominant effects, the multicopy nature of *rrs* substantially reduces both the detectability and predictive power of *rrs*-based intrinsic resistance.

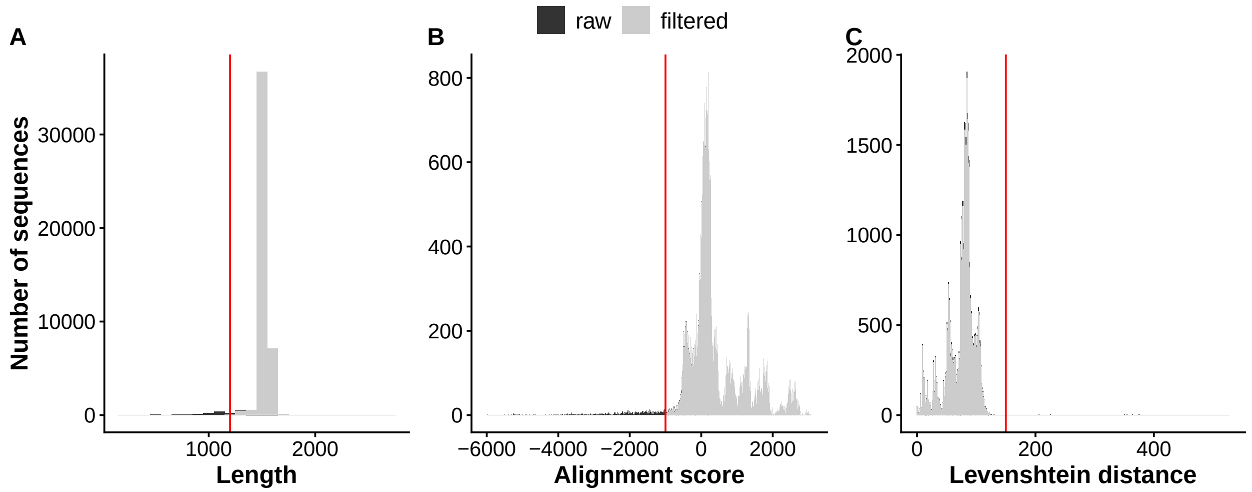


**Figure S7.** Properties of *rrs* sequences extracted from genomes downloaded from NCBI. The plots show the distribution of key sequence properties before (black) and after (grey) filtering. A) Distribution of sequence lengths, with a vertical red line indicating the minimum length threshold of 1200 bp. B) Alignment scores of each sequence against the *E. coli rrs* reference sequence, with a cut-off of −1000. C) Levenshtein distance of the entire sequence relative to the *E. coli* reference, with a red line indicating the maximum threshold of 150 used for filtering.

**Alt text:** Histograms of *rrs* sequence properties before and after quality filtering, subfigures A, B, and C showing distributions of sequence length, alignment score, and Levenshtein distance relative to the *E. coli* reference, with filtering thresholds indicated.

**Table S1.** Taxonomic summary of species predicted to exhibit intrinsic STR resistance from *rrs* analysis. The total number of *rrs* gene copies identified per species and the number of copies carrying mutations associated with intrinsic resistance are reported.

| **Species** | ***rrs* gene copies** | ***rrs* gene copies with resistance variants** | **Class** |
|---|---|---|---|
| *Arsenophonus endosymbiont of Bemisia tabaci* | 7 | 2 | Gammaproteobacteria |
| *Dictyobacter kobayashii* | 9 | 1 | Ktedonobacteria |
| *Moritella marina* | 17 | 4 | Gammaproteobacteria |
| *Pectobacterium parvum* | 7 | 1 | Gammaproteobacteria |
| *Psychrosphaera algicola* | 5 | 3 | Gammaproteobacteria |
| *Spiroplasma taiwanense* | 1 | 1 | Bacilli |
| *Vibrio aquimaris* | 9 | 1 | Gammaproteobacteria |
| *Zobellia laminariae* | 3 | 1 | Bacteroidia |

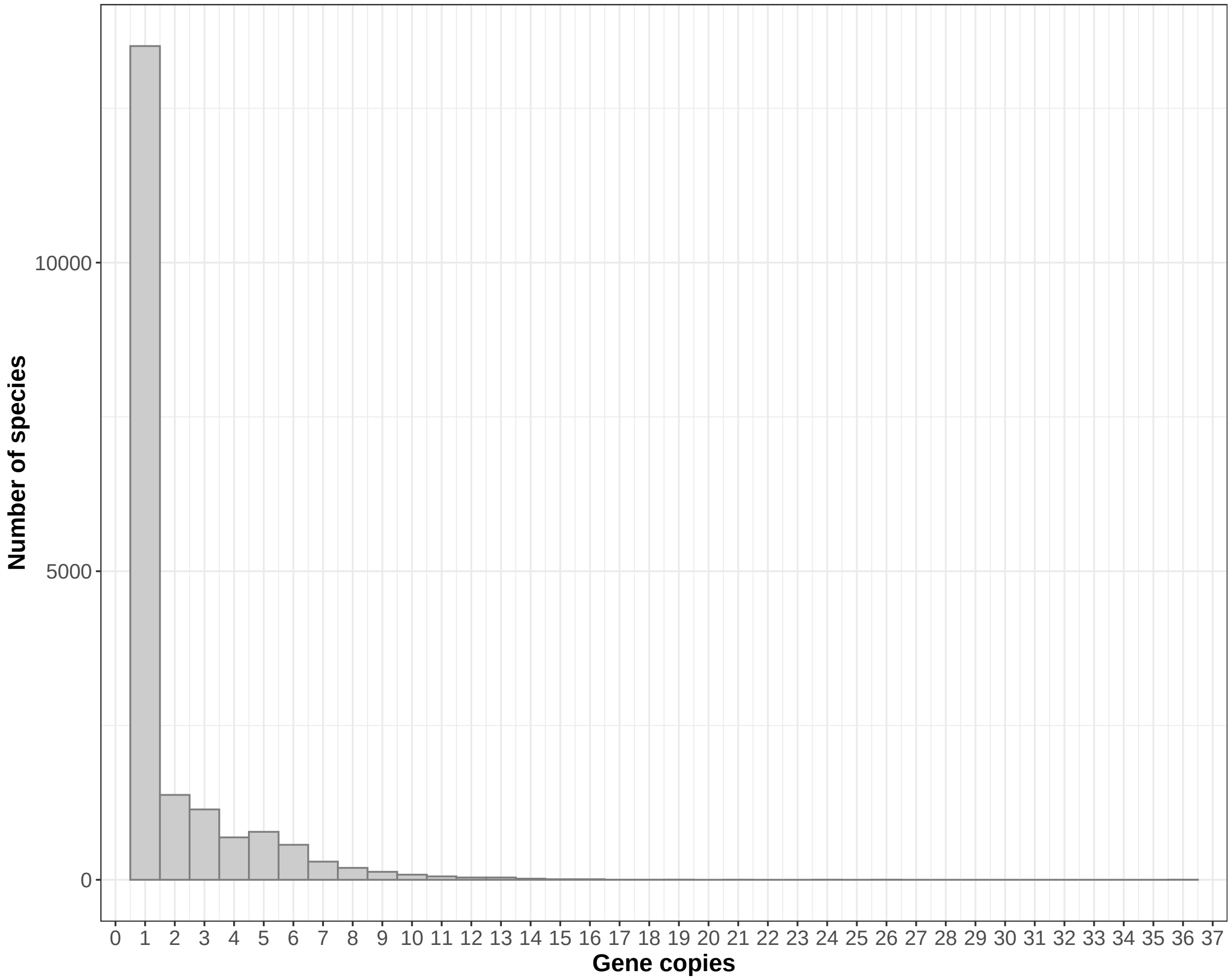


**Figure S8.** Distribution of *rrs* gene copy number. The histogram shows the number of species observed for each *rrs* copy number.

**Alt text:** Histogram showing the distribution of *rrs* gene copy number across species, with most species carrying a single copy and copy numbers ranging up to 37.
